# Modulation of early level EEG signatures by distributed facial emotion cues

**DOI:** 10.1101/2021.03.03.433729

**Authors:** Diana Costa, Camila Dias, Teresa Sousa, João Estiveira, João Castelhano, Verónica Figueiredo, Andreia Pereira, Miguel Castelo-Branco

## Abstract

Face perception plays an important role in our daily social interactions, as it is essential to recognize emotions. The N170 Event Related Potential (ERP) component has been widely identified as a major face-sensitive neuronal marker. However, despite extensive investigations conducted to examine this electroencephalographic pattern, there is yet no agreement regarding its sensitivity to the content of facial expressions.

Here, we aim to clarify the EEG signatures of the recognition of facial expressions by investigating ERP components that we hypothesize to be associated with this cognitive process. We asked the question whether the recognition of facial expressions is encoded by the N170 as weel as at the level of P100 and P250. In order to test this hypothesis, we analysed differences in amplitudes and latencies for the three ERPs, in a sample of 20 participants. A visual paradigm requiring explicit recognition of happy, sad and neutral faces was used. The facial cues were explicitly controlled to vary only regarding mouth and eye components. We found that non neutral emotion expressions elicit a response difference in the amplitude of N170 and P250. In contrast with the P100, there by excluding a role for low level factors.

Our study brings new light to the controversy whether emotional face expressions modulate early visual response components, which have been often analysed apart. The results support the tenet that neutral and emotional faces evoke distinct N170 patterns, but go further by revealing that this is also true for P250, unlike the P100.

## 1. Introduction

The dynamics of face perception involve several brain regions and circuits responsible for the early and high level visual processing of faces(1). Studies using functional magnetic resonance imaging (fMRI) can identify the brain regions underlying these processes but cannot capture their temporal properties. Thus, the dynamics of face perception neuronal mechanisms have been extensively studied based on electroencephalographic (EEG) signals (2).

Social attention has been used as a synonym for non-verbal social communication behaviors. Of socially relevant stimuli, faces and gazes are the two most important elements triggering this cognitive process (3) (4). Categorization facial expressions, implies allocation of selective attention which may also modulate the neural response differences between the categories of emotions. When the participants perform implicit tasks more directed to the aspect emotional expression, there seems to be a lower probability of finding differences between the categories of emotions, namely for the N170 component (5).

Facial expressions elicit robust neuronal responses that can be measured through event-related potentials (ERP) analysis. The lateral occipitotemporal face-sensitive N170 component is generally used as a major neuronal marker of face recognition. However, the literature is not unanimous about how N170 is affected by the emotional content of facial expressions(6,7). Some authors suggest N170 occurs as a result of early automatic structural encoding of faces, which occurs before a comparison of these structural descriptions with representations stored in memory (8,9). On the other hand, others challenge the view that structural encoding is temporally distinct from emotion processing and defend that the N170 can be modulated by emotional expressions, as shown by larger amplitude and longer latency (10–12).

Previous findings also suggest the involvement of P100 in face-specific visual processing (13,14). However, this posterior component often also assumed to have an extrastriate contribution is believed to reflect low-level visual features processing and, therefore, the P100 role in face recognition remains an open question (5). Another potentially relevant ERP is the P250. Like the commonly measured N170, the P250 is also maximal at occipito-temporal sites. There is evidence for P250 modulation during face perception (15,16), but this ERP component has been often related to higher-level nonspecific aspects of face processing (17,18).

According to Puce et al. (19), these ERPs can be grouped in a robust positive-negative-positive (P100-N170-P250) ERP complex, serving as neuroelectric markers in the investigation of the visual processes involved in the recognition of facial expressions. There is also evidence that both P100 and N170 are also involved in generating differencial responses to neutral vs. emotional expressions, similarly regions such as the amygdala (20).

Importantly, temporal information of face perception provided by EEG may be coupled with source localization techniques to identify their cortical origin. The inferior occipital gyrus (IOG), the lateral portion of the fusiform gyrus (FG), and the superior temporal sulcus (STS) have been described as the core system of face perception. IOG and FG are believed to mediate the encoding of faces, while STS is involved in the perception of social signals derived from faces, such as the direction of gaze and emotional expressions (21,22). However,the relationship between N170 and signal originating from these face-selective brain regions remains under discussion. Recent findings suggest the FG to be a main neuronal contributor to the N170 ERP component (2).

Here, we aim to contribute to this debate concerning the temporal dynamics and the electrophysiological nature of face perception. First, we seek to understand if we can objectively distinguish the emotional content of the facial expressions based on not only the N170 ERP component but also on direct comparisons with the P100 and P250. Then, we want to address whether the investigation of the neuronal sources underlying these temporal patterns can shed light on this debate.

To achieve this goal, we used an experimental paradigm in which the participants had to identify the facial expression presented on the screen to generate an action. The study was organized in two experiments, the only difference being the inclusion of a neutral face in the instruction phase, in experiment 2. Our results show that ERPs N170 and P250 are both modulated by facial expressions (happy and sad). Differences were only found between the conditions expressions – neutral faces but not between the expressions per se.

## 2. Materials and Methods

### 2.1 Experimental Design

Two experiments were performed, each on a different group of subjects. Below we describe together the attributes of the participants, as well as data acquisition, EEG processing and data analysis, since the information is similar between experimets. Separately we present the stimulation paradigms and the results of the two experiments.

The use of the paradigm described below required participants to recognize facial expressions which was informative for a subsequent action related to error monitoring neural processes (not reported in this paper). For this, two experiments were performed, the main tak objective is the recognition and distinction between facial expressions (happy, sad and neutral) by the participant. The two experiments differ in the existence or not of a neutral face in the initial period (Experiment 1) or in the instruction phase (Experiment 2) which allowed to control for the role of an explicit instruction for the neutral stimulus.

### 2.2 Participants

In total, 40 adult with normal or corrected-to-normal vision and no medical or psychological disorders were, included in this study: In the experiment one, 20 individuals were included (nine females), mostly right-handed (19), and aged between 20 and 36 years (26.8 ± 4.514); in the experiment two, 20 individuals were included (11 females), (all right-handed), and aged between 20 to 33 years (25.7 ± 4.193), of these subjects, 5 participated in both experiments.

Written informed consent was obtained from all participants. The study was approved by the ethics committee of the Faculty of Medicine from the University of Coimbra and was conducted in accordance with the declaration of Helsinki.

### 2.3 Stimuli

#### 2.3.1 Experiment 1

A standard set of colour photographs consisting of a male, each depicting happy, sad, and neutral faces, obtained from the Radboud Faces Database (23), were presented on a 17-inch monitor situated 60 cm away from the subject. The facial expression images had a mean luminance of 7.67×10^1^ cd/m^2^ (with screen luminace ranging from 2.44×10^1^ cd/m^2^ to 1.76×10^2^ cd/m^2^).

Neutral, happy, and sad faces (height 4.55° and width 4.90°) were presented to the participants in a pseudo-randomized order (Figure 1A). A go/no-go task, based on facial expression recognition, was used to guarantee implicit expression processing (Figure 1B).

**Fig 1.**
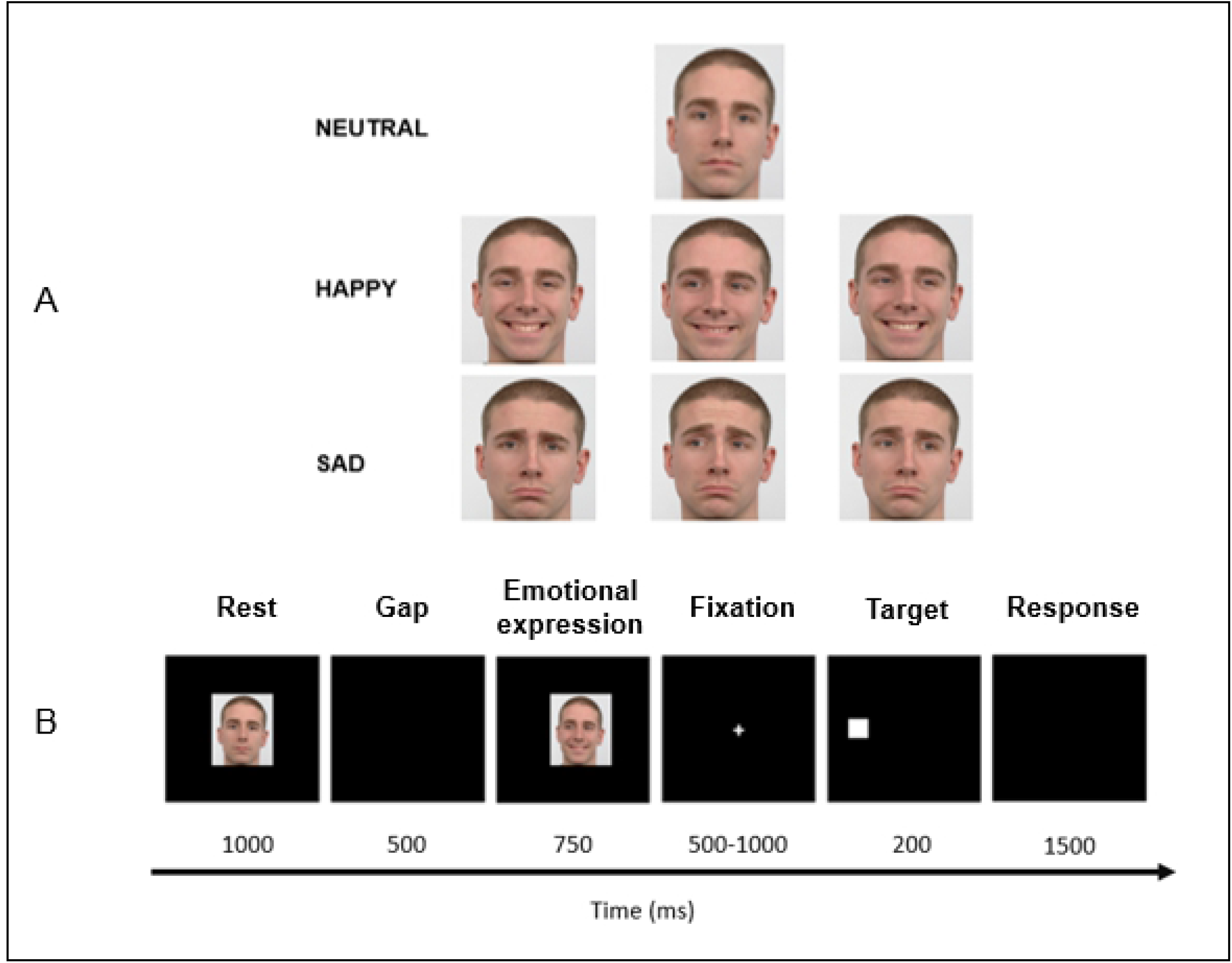
Overview of the facial stimuli paradigm used in experiment 1. Neutral, happy, and sad faces (A) were used as cues in a go/no-go paradigm (B) and promoted attentive expression processing. The sequence between face presentation and participants’ response is also illustrated.

The stimulation paradigm included six conditions: rest, gap, emotional expression, fixation, target, and response. Initially, a neutral face was displayed for 1000 ms (rest period), followed by a gap period of 500 ms. Then, the instruction for the go/no-go task is presented during 750 ms. Here, a happy, or sad face was presented. The expression type cued the subsequent action, and thereby required attentive processing.A happy face with a gaze means go and perform a saccade in the same gaze direction; a sad face with a gaze means go and perform a saccade in the opposite gaze direction; a happy or sad face without a gaze means to no-go. After a fixation period (500 - 1000 ms) and a target shown for 200 ms in the same direction as the gaze (height and width 0.72°), a black background appeared for 1500 ms, where the participants performed the task.

#### 2.3.2 Experiment 2

A standard set of colour photographs consisting of a male, each depicting happy, sad, and neutral faces, obtained from Radbound Faces Database (23), were presented on a () monitor situated 60 cm away from the subject. Neutral, happy, and sad faces (height 9.02° and width) were presented to the participants in a pseudo-randomized order (Figure 1A). A go/no-go task, based on facial expression recognition, was used to guarantee implicit expression processing (Figure 1B).

As in experiment 1, this second paradigm also includes six conditions: rest (neutral faces), gap 1, social expression, gap 2, action instruction, and action. Initially, a neutral face was displayed for 1000 ms (rest period), followed by a gap period of 200 or 500 ms. Then, the instruction (social expression) for the go/no-go task is presented during 350 ms. Here, a happy, sad or neutral face was presented. It is in this condition that the biggest difference between the two experiences, in experiment one we only had happy and sad as social expression. A happy face with a gaze means go and perform a saccade or button press in the same gaze direction; a sad face with a gaze means go and perform a saccade or button press in the opposite gaze direction; a happy, sad or neutral face without a gaze means to no-go. After a gap 2 period (200 ms) and an action instruction been shown for 350 ms (target diamond (♦) – height and weidth 2.517°) or button press (target: square (□), – height and weidth 1.819), a black background appeared for 1000 ms, where the participants performed the task.

### 2.4 Experimental setup and data recording

EEG data were recorded using a 64 electrodes cap (Compumedics Quick cap; NeuroScan, USA). The scalp of the participants was first cleaned using abrasive gel and then the electrodes cap was placed on their head according to the international 10–20 standard system.

Electrooculogram (EOG) data were recorded via two pairs of additional electrodes, placed above and below the left eye and in the external corner of both eyes. The reference electrode was located between Cz and CPz The impedance of the electrodes was kept under 20 kΩ during the recordings. The electrodes were connected directly to the SynAmps 2 amplifier system (Compumedics NeuroScan, Texas, USA) and sampled at 1000Hz. Data were recorded using the Curry Neuroimage 7.08 (NeuroScan, USA). For each paradigm, the participants were informed about the respective task. The total duration of the experimental procedure including the preparation procedures took around 60 min.

The eyetracking (ET) data acquisition started with the calibration of the eye-tracker. The data were recorded at 120Hz in a tower-mounted high accuracy (0.25° – 0.5°) monocular eye-tracker (iView X™ Hi-Speed 1250Hz, SMI – SensoMotoric Instruments, Teltow, Germany).

### 2.5 EEG processing

In both experiments, we used MATLAB home-made script (R2018b) and the EEGLAB toolbox functions (version 2) for EEG signal preprocessing and analysis.

The EEG data were downsampled to 500 Hz and filtered between 0.5 and 45 Hz. Noisy channels were removed and the electrodes were re-referenced to the average of all EEG (excluding EOG) channels. The data were segmented into epochs of 1700 ms length in experiment 1 and 1200 ms in experiment 2, with a 200 ms pre-instruction baseline. Epochs were visually inspected, and noisy trials were removed. Ocular, muscular and cardiac artifacts were removed from all EEG channels based on the independent components analysis (ICA) (24). The noisy channels previously removed were interpolated (spherical interpolation).

After pre-processing, in experiment 1 there were on average 150, 143 and 143 epochs per participant for happy, sad and neutal faces respectively, in experiment 2 there were on average 156, 158 and 155 epochs for happy, sad and neutral faces respectively.

### 2.5 Data Analysis

#### 2.5.1 Event Related Potentials (ERPs) identification

The N170 ERP component has been described as a negative peak around 170 ms after the stimulus beginning, particularly incident at the parieto-occipital electrodes (25,26), whereas, P100 and P250 are ERP positive components. Their topology has also been described in the lateral parieto-occipital cortex as all these ERP components related to rapid processes of selective attention (27,28) . Therefore, we started by selecting the groups of electrodes where these ERPs were most prominent. This group of electrodes was T8, CP6, TP8, P8 (right hemisphere) and TP7, P7, P5, PO7 (left hemisphere) since they showed the most significant difference between the three expressions, in N170 component(p <0.01 with Bonferroni correction). ERPs were first identified through the maximum (P100 and P250) and minimum (N170) peaks of each trial in the following P100 and N170 a time range of 50 to 200 ms and P250 a time renge of 200 to 300 ms, in both experimets. Then, ERPs components for each experimental condition of interest (happy, sad, and neutral faces) were estimated per participant as the average of the identified peaks. Finally, N170, P100 and P250 were statistically compared for the three different expressions types in order to test for differences in amplitude, latency and lateralization.

#### 2.5.2 Source Estimation

Source analysis was conducted using the sLORETA software (29) (pascual-marqui). The procedure included exporting from MATLAB the preprocessed single-trial epochs, importing them into sLORETA software, averaging them (per subject and expression) and converting them to the source space. Each participant electrode location was co-registered with the realistic anatomical model using landmarks and standard electrode’s position. The sLORETA uses a three-shell spherical head model registered to the digitized (30) atlas (Brain Imaging Centre, Montreal Neurological Institute). The solution space is restricted to cortical gray matter and the hippocampus.The sLoreta yields images of standardized current source density of a total of 6430 voxels at 5 mm spatial resolution under these neuroanatomical constraints. Briefly, subject time and valence specific mean ERPs were transformed into the sLoreta domain and averaged across subjects within each experimental group. Localization of power was then determined for time windows 100 ms, 170 ms and 250 ms.

#### 2.5.3 Statistical Analysis

IBM SPSS statistics 25 was used for statistical analysis. The maximum peak to peak amplitude differences between the face expressions (happy and sad) and neutral faces was tested by Pairwise methods:-Anova repeated measures with happy, sad and neutral faces as a factor, was used to examine the overall effects for different amplitudes, latencies, and peaks. This analysis was performed for both hemispheres (right and left). A P value of <0.05 was significant. The same statistical analyses were performed for the three ERPs (P100, N170, P250).

## 3. Results

### 3.1 ERPs amplitude and latency

In the analysis of the ERPs, the amplitude and latency of its main components, P100, N170 and P250 were estimated per participant and condition and compared. The analyses were performed per hemisphere to allow testing for lateralization effects. The results are, presented per experiment. For these analyses two clusters of channels were used, T8, TP8, P6, P8, (right hemisphere) and TP7, P7, P5, PO7, (left hemisphere).

### 3.2 Experiment 1

#### 3.2.1 P100

When comparing EEG responses to happy, sad, and neutral faces at the level of P100 we found neither differences in amplitude nor latency. The grand average ERP waveforms are illustrated in Figure 3, whereas the statistical results are summarized in Figure 4.

**Fig 2.**
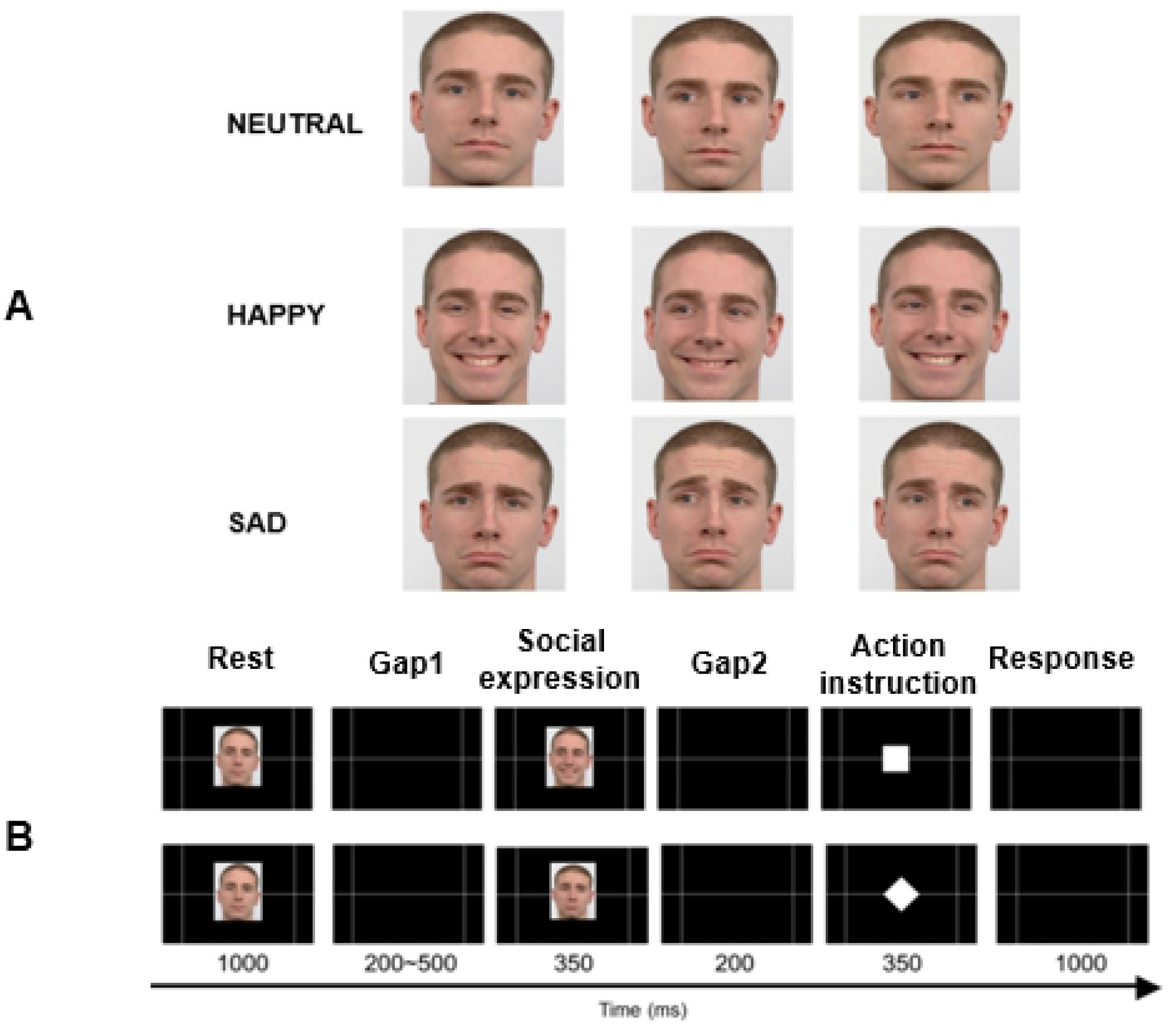
Overview of the facial stimuli paradigm used in experiement 2. Neutral, happy, and sad faces (A) were used as cues in a go/no-go paradigm (B) and allowed to study the implicit expression processing. The sequence between face presentation and participants’ response is also illustrated.

**Fig 3.**
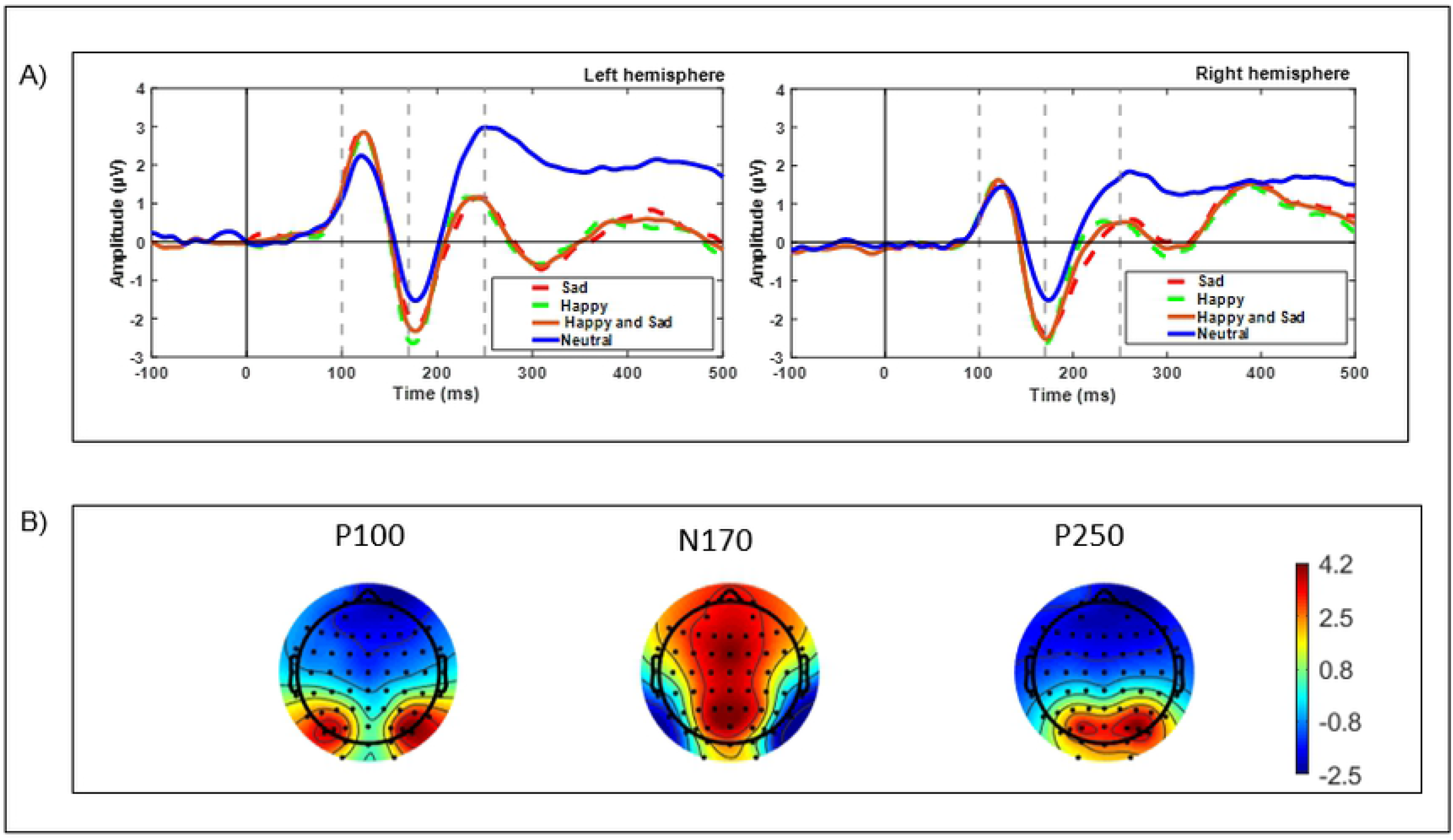
Grand-average ERP waveforms from the left and right hemispheres for happy and sad expressions, and neutral faces. A) The 100 ms, 170 ms, and 250 ms are highlighted with a dashed line. The results presented are relative to the channel clusters TP7 P7, P5, PO7 at the left hemisphere, and T8, CP6, TP8, P8 at the right hemisphere. B) Scalp topographic voltage maps for the emotions (happy and sad) stimulus condition. Maps reflect the activity profile at the following post-stimulus onset latencies: P100 (100 ms) N170 (170 ms), and P250 (250 ms).

**Fig 4.**
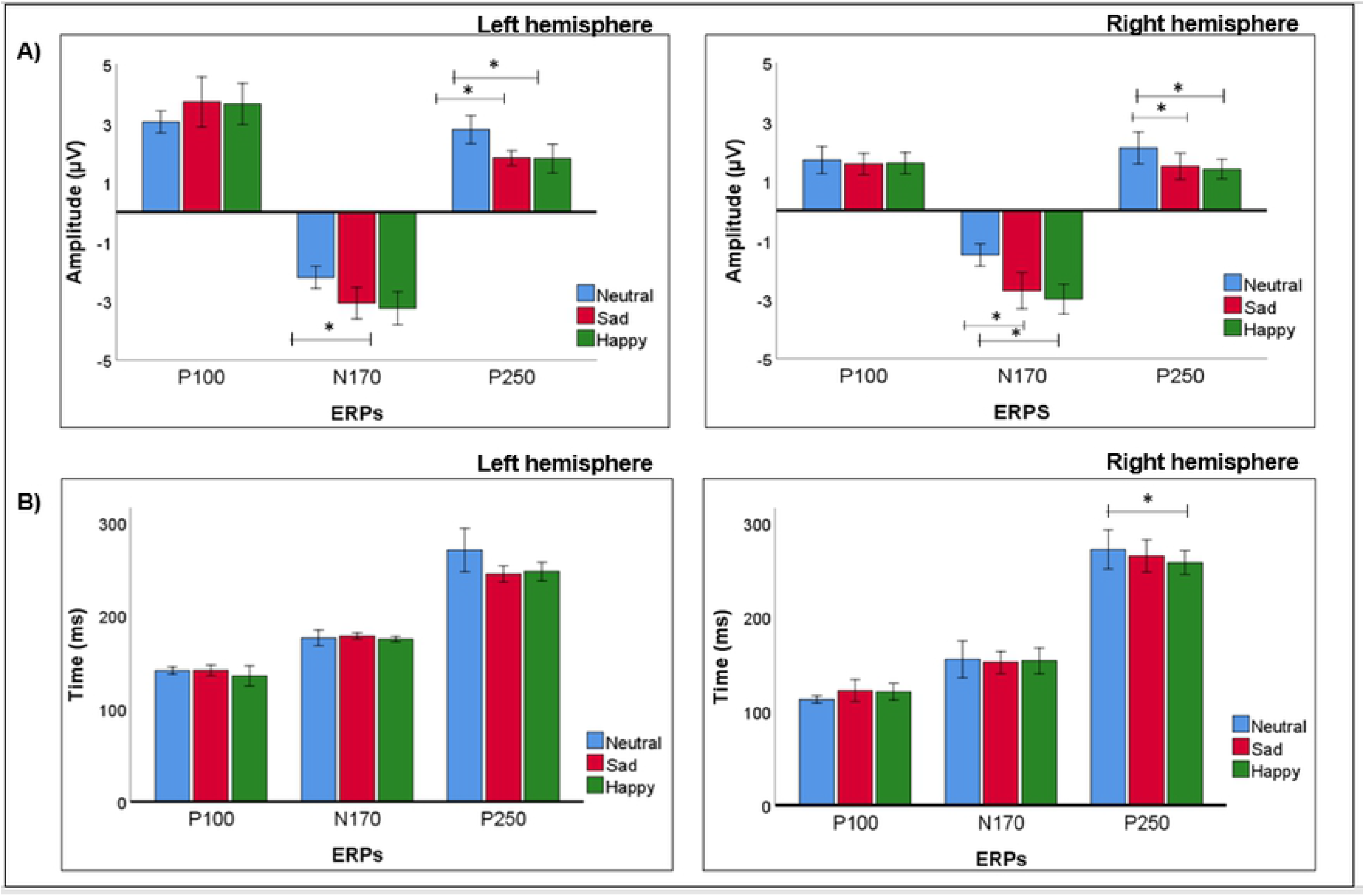
Summary of amplitude and latency analyses for the ERP components P100, N170, and P250 when comparing brain responses during emotional and neutral faces recognition (Experiment 1). A) We found significant differences in all ERPs between neutral faces and expressions (happy and sad), except for P100 in both hemispheres. B) We only found significant latency differences for the P250, in the right hemisphere, between neutral faces and happy faces. Error bars depict the standard error of the mean.

In the right hemisphere the P100 average amplitude was 1.959±0.308 µV, 1.952±0.340 µV, and 1.929±0.277 µV for happy, sad and neutral conditions respectively. Peaks of latencies were recorded,123 ms after happy and sad stimuli and 122 ms after neutral stimuli. In the left hemisphere, were recorded amplitudes of 3.809±0.601 µV for happy faces, 3.994±0.656 µV for sad faces and 3.311±0.563 µV for neutral faces. Latencies in this hemisphere were recorded at 122 ms after happy and 123 ms after sad and neutral stimuli respectively.

#### 3.2.2 N170

When comparing the three conditions, at the level of N170 we found differences in the amplitude between neutral-sad in the right hemisphere and between neutral-happy in both hemispheres. The grand average ERP waveforms are illustrated in Figure 3, whereas the statistical results are summarized in Figure 4.

In the right hemisphere the N170 average amplitude was -2.976±0.330 µV, -2.769±0.339 µV, and -1.727±0.332 µV for happy, sad and neutral conditions respectively, significant differences were found between neutral - sad,1.032±0.358 µV, p = 0.03 and neutral – happy, 1.239±0.4 µV, p = 0.02, F(2,18)=4.560, p= 0.025. Latencies were recorded at 172 ms after happy stimuli, 171 ms after sad and 174 ms after neutral stimuli. In the left hemisphere the average amplitude was - 3.461±0.420 µV, -3.213±0.373 µV and -2.573±0.436 µV for happy, sad and neutral conditions, significant differences were found between neutral – happy, 0.888±0.290 µV, p = 0.02, F(2,18)=4.640, p=0.024. Latencies were recorded at 175 ms after happy stimuli, and 176 ms after sad and neutral stimuli.

#### 3.2.3 P250

When comparing the three conditions at the level of P250 we found differences in the amplitude between neutral-sad and neutral-happy in both hemispheres and in the latency neutral and happy in right hemisphere. The grand average ERP waveforms are illustrated in Figure 3, whereas the statistical results are summarized in Figure 4.

In the right hemisphere, the P250 average amplitude was 1.290±0.379 µV, 1.237±0.387 µV, and 2.260±0.577 µV for happy, sad and neutral faces. We found significant differences between neutral – sad, 1.024±0.324 µV, p = 0.015 and neutral – happy, 0.970±0.318 µV, p = 0.02, F(2,18)=4.879, p=0.02. Latencies were recorded at 240 ms after happy stimuli, 246 ms after sad and 253 ms after neutral stimuli, significantive differences were found between neutral – happy, 13.8±4.502 ms, p=0.019, F(2,18)= 6.185, p=0.009. In left hemisphere, significant differences were recorded at amplitudes of 1.812±0.428 µV for happy, 1.804±0.376 µV for sad and 2.949±0.505 µV for neutral faces, found between neutral – sad,1.145±0.349 µV, p = 0.012 and neutral – happy, 1.136±0.321 µV,p = 0.007, F(2,18)=6.094, p=0.01. Latencies were recorded at 236 ms after happy stimuli, 235 ms after sad stimuli and 243 ms after neutral stimuli.

### 3.3 Experiment 2

#### 3.3.1 P100

When comparing EEG responses to happy, sad, and neutral faces at the level of P100 we found neither differences in amplitude nor latency. The grand average ERP waveforms are illustrated in Figure 5, whereas the statistical results are summarized in Figure 6.

**Fig 5.**
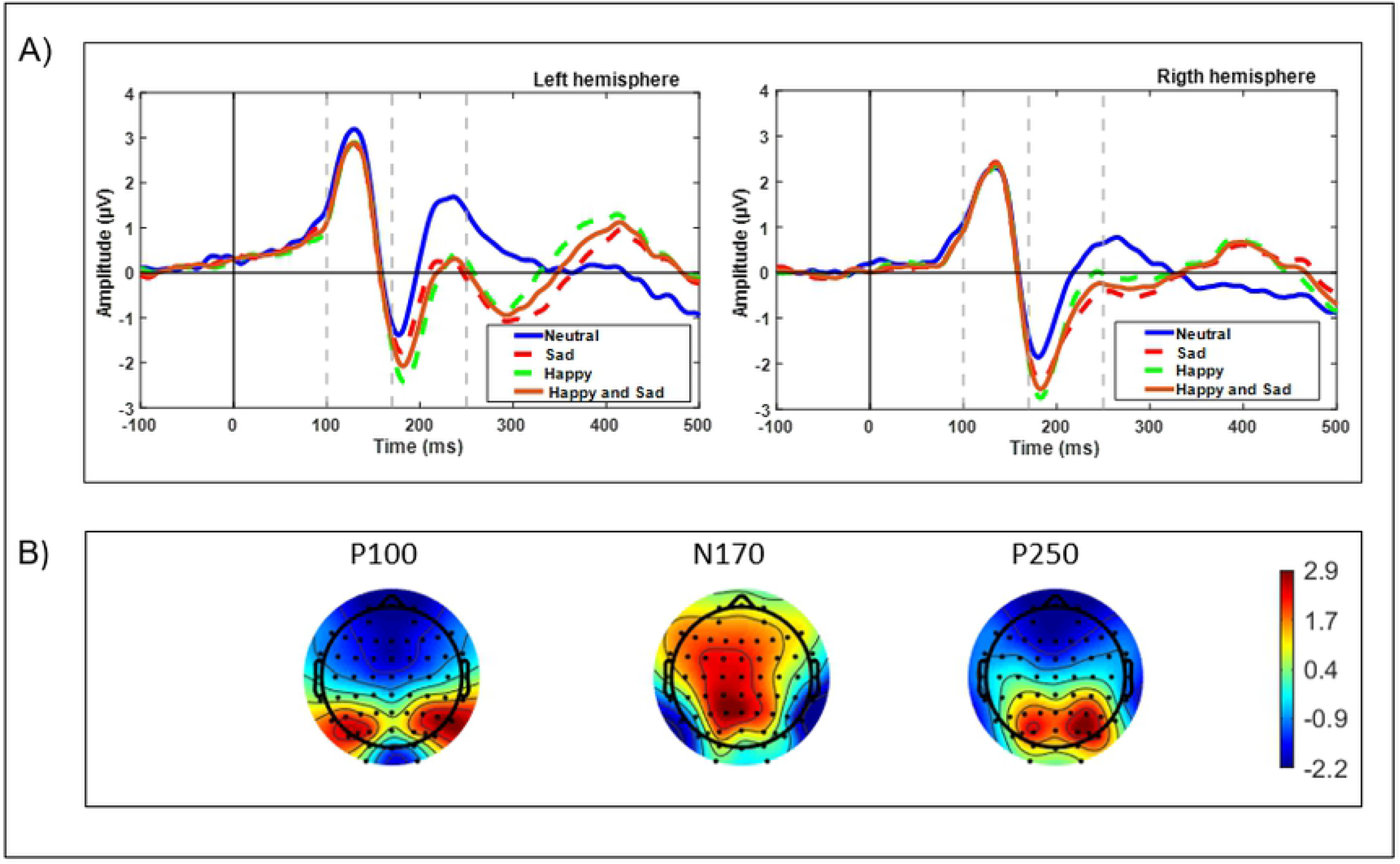
Grand-average ERP waveforms from the left and right hemispheres for happy and sad expressions, and neutral faces. The 100 ms, 170 ms, and 250 ms are highlighted with a dashed line. The results presented are relative to the channel clusters TP7 P7, P5, PO7 at the left hemisphere, and T8, CP6, TP8, P8 at the right hemisphere. B) Scalp topographic voltage maps for the emotions (happy and sad) stimulus condition. Maps reflect the activity profile at the following post-stimulus onset latencies: P100 (100 ms) N170 (170 ms), and P250 (250 ms).

**Fig 6.**
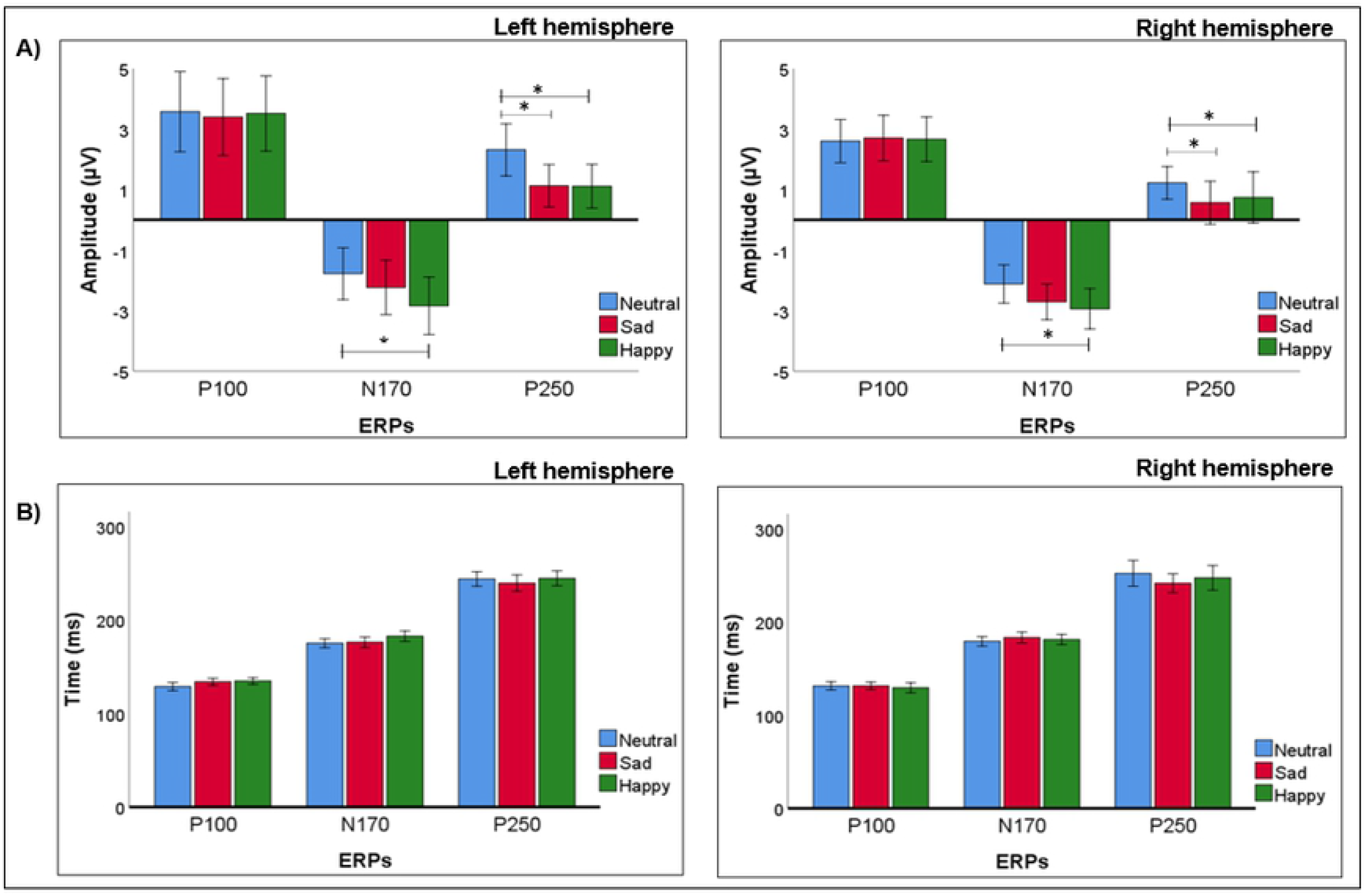
Summary of amplitude and latency analyses for the ERP componets P100, N170, and P250 when comparing brain responses during emotional and neutral faces recognition (Experiment 2). A) We found significant differences in all ERPs between neutral faces and expressions (happy and sad), except for the P100 in both hemispheres. B) We did not find significant latency differences for ERPs in neither of the hemispheres. Error bars depict the standard error of the mean.

In the right hemisphere the P100 average amplitude was 3.229±0.651 µV, 3.367±0.602 µV and 3.359±0.581 µV for happy, sad and neutral conditions respectively. Latencies were recorded at 129 ms after happy stimuli, 130 ms after sad and 132 ms after neutral stimuli. In the left hemisphere, P100 average amplitude was 4.488±0.877 µV, 4.611±0.946 µV and 5.007±0.958 µV for happy, sad and neutral faces. Latencies were recorded at 131 ms after happy stimuli and 133 ms after sad and neural stimulis.

#### 3.3.2. N170

When comparing EEG responses to happy, sad, and neutral faces at the level of N170 we found differences in the amplitude between neutral and sad faces in both hemispheres. The grand average ERP waveforms are illustrated in Figure 5, whereas the statistical results are summarized in Figure 6.

In the right hemisphere the N170 amplitude average was -3.067±0.457 µV, -2.881±0.410 µV, and -2.153±0.309 µV for happy, sad and neutral conditions respectively, significative differences were found between between neutral – happy, 0.728±0.246 µV p=0.047, F(2,18)=8.152, p=0.003. Latencies were recorded at 183 ms after happy and sad stimulis and 180 ms after neutral stimuli. In the left hemisphere, amplitude average for N170 was -2.906±0.482 µV, -2.500±0.619 µV and -1.737±0.615 µV for happy, sad and neutral conditions, significative differences were found between neutral - happy, 0.763±0.147 µV, p = 0.002, F(2,18)=11.774, p=0.001. Latencies were recorded at 179 ms after happy stimuli and 176 ms after sad and neutral stimulis.

#### 3.3.3 P250

When comparing EEG responses to happy, sad, and neutral faces at the level of P250 we found differences in the amplitude between neutral and happy, and neutral and sad in both hemispheres. The grand average ERP waveforms are illustrated in Figure 5, whereas the statistical results are summarized in Figure 6.

In the right hemisphere the P250 amplitude average was 1.127±0.397 µV, 0.923±0.599 µV and 2.098±0.452 µV for happy, sad and neutral conditions respectively,significative differences between neutral – sad, 1.175±0.343 µV, p = 0.023 and neutral – happy µV, p = 0.028, F(2,18)=11.850, p=0.001. Latencies were recorded at 256 ms, 252 and 259 ms after happy, sad and neutral stimulis respectively. In the left hemisphere, amplitude average for P250 was 0.989±0.558 µV, 1.416±0.583 µV and 2.686±0.691 µV for happy, sad and neutral respectively, significative differences were found between neutral – sad, 1.271±0.270 µV, p = 0.003 and neutral – happy, 1.697±0.472 µV, p = 0.017, F(2,18)=18.008, p=0.0002. Latencies were recorded at 240 ms, 243 ms and 246 ms after happy, sad and neutral stimulis.

## 4. Source Estimation for the ERPs components modulated by social attention

The results of the sLoreta for ERPs modulated by emotional expressions (happy and sad) for 170 ms and 250 ms post-stimulus presentation onset are depicted in fig.7.

**Fig 7.**
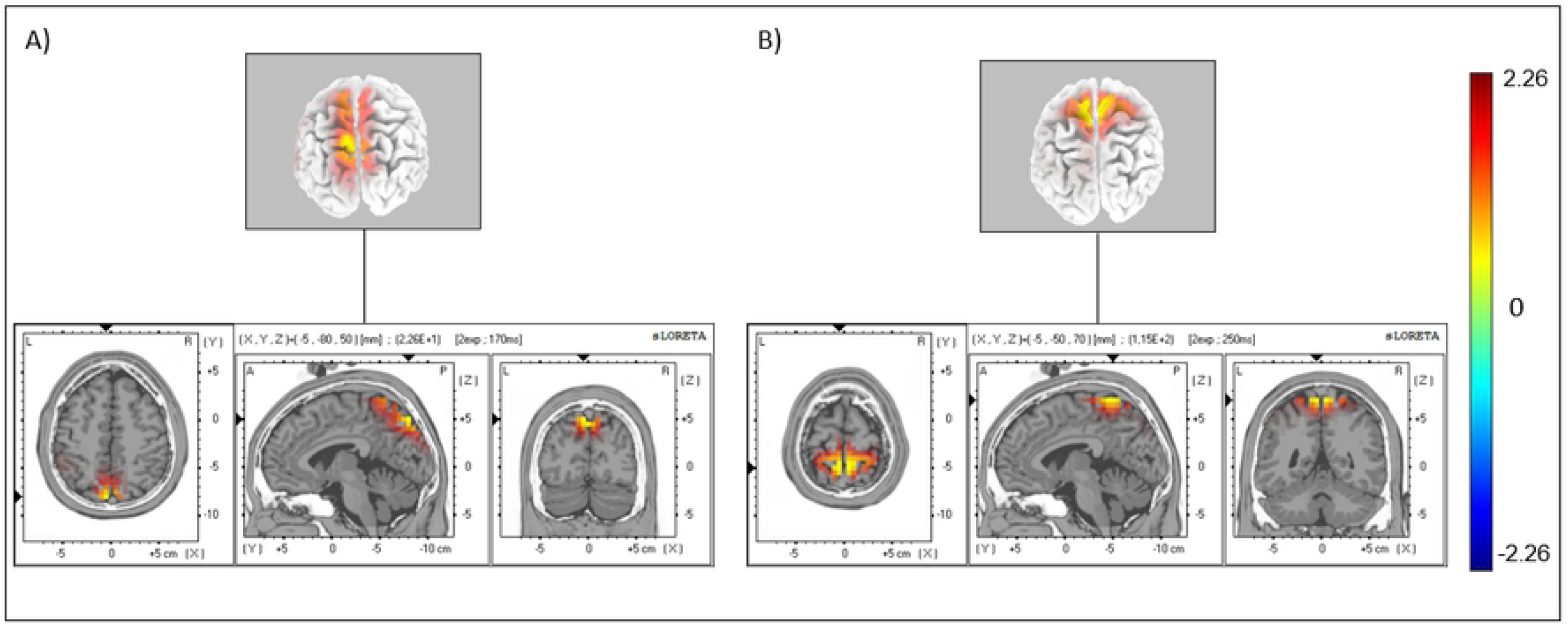
sLORETA representation of significant correlations of emotional expressions for N170 and P250. A) N170 source located in the parietal lobe and precuneus B) P250 source located in the parietal lobe, postcentral gyrus.

Sources estimation for expression at 170 ms (N170) was found in the precuneus, BA 7 (MNI coordinates: x= -5, y= -80, z= 50) and at 250 ms (P250) was found in the postcentral gyrus, BA 5 (MNI coordinates: x= -5, y=-50, z= 70), significant at P = 0.0002, two-tailed t test.

## 5. Discussion

In this study, we investigated the temporal dynamics of the neural mechanisms underlying face recognition, following the hypothesis that not only the N170 but also other related components as P100 and P250 are modulated by the emotional content of facial stimuli. To this end, we used an implicit face recognition task, while controlling for selective attention. We tested for amplitude and latency differences between neutral and emotional expressions, particularly happy and sad faces. Moreover, we aimed at exploring the neural sources of social attention.

N170 is a well-established ERP component of face processing (4) . However, there is no consensus regarding N170 selectivty for the content of facial expressions. Some studies support that the presence of N170 amplitude dependence on emotional facial expressions (31,32), whereas others suggest that this ERP is modulated by more low level features of facial stimuli but does not discriminate between expressions (9,33). Our results contribute to this debate by showing the presence of an effect when attention is controlled for.

Our results show indeed that there is a clear difference in electrophysiological responses between expressions (happy and sad) and neutral faces. In both experiments, a similar pattern was observed, we did see a desponse difference between facial expressions from neutral faces (except for the P100), with no significant differences between sad and happy faces, in both experiments. These results are consistent with previous studies in which the N170 was affected by emotion expression, specifically happy and fearful faces had larger amplitude than neutral faces (34), but did not discriminate between different emotions (35). N170 has been proposed to represent an early configural stages of face processing, which may reflect an activation related to the structural coding of faces, and from this point of view response levels might be expected to be relatively invariant to emotional details, in facial expressions.However, Luo, et al., 2010 (33) proposed that ERPs between the 150 to 300 ms time period constitute a “second stage” of expression processing that is already sensitive to emotionality in general (compared to a neutral expression); arguing that the differentiation between specific expressions / emotions occurs only in a “third stage” that starts after 300ms.

The earliest stage of visual processing, P100 is associated with processing sensory characteristics of a visual stimulus, there by unlikely to be modifiable by familiarity and expressions (36). However, there are studies that suggest that differentiation between expressions can occur, such as fear and anger or happiness and anger (37,38).

According to our results, no significant difference was identified between the P100 amplitudes in the three facial expressions (happy, sad and neutral),as such we argue that this ERP component is not modulated by expressions. P100 can also be modulated by attention, so the increase in amplitude in prior studies may be due not only to expressions but also to increased effects of attention (13). Under controlled attention, (futher optimized in experiment 2), we found no significant effect of the presence of emotion cues, concerning the P100. For comparison, there is corroborating evidence that as early as the P100 component, negative and positive expressions, such as fear, sadness or happiness, cannot be differentiated (32,37). We did not observe P100 amplitude significant difference emotions types (happy and sad), or between both emotional faces types and the neutral faces. These results are in agreement with other authors, such as (25,39).

P250 is the last ERP that forms the P100-N170-P250 complex, according to previous studies like (40) this ERP is responsive to emotional expressions, amplitudes are augmented for emotional expressions compared to neutral.

Our results show, in both experiments, an amplitude increase (more negative) in happy and sad facial expressions for the P250. Unlike the previous ERPs (P100, N170), the neutral faces were the ones with the highest amplitudes, with significant differences with the happy and sad amplitude.

In this work, latency was only significant in the right hemisphere of the first experiment, in ERP P250 between neutral and happy faces. These results do agree with previous studies (41), who found latency change for expressions when compared to neutral stimuli, justifying that this difference reflected the impact voluntary attention, which is also consistent with the conditions of experiment 1.

Both in experiment 1 and 2, differences were only found between the conditions expressions – neutral faces and not between the emotional expression types happy – sad, showing that the differences between both experiments did not change the general patterns of results.

Our source analysis allowed to investigate the spatical location of the signals studied, as such, to analyse the sources of facial expressions and to compare results from previous studies . It wasexpected that the location of the active sources should be found in the parieto-occipital region, because this region contributes mainly to the processing of facial expressions (42). N170 and P250 were modulated by the expressions happy and sad, so we went to analyse where the sources of each one are located, and confirmed that they match The source of N170 was identified in the precuneus. This region is involved in visuospatial processing, directing attention in space and is one of the core regions of the perspective taking network, both from a cognitive and affective point of view (43,44). The source of the P250 was located in the postcentral gyrus that belongs to the superior parietal lobe (SPL) which are major areas in the attention central system(45). The precuneus is also selectively connected to the SPL. Converging evidence then suggests that the SPL and the precuneus cooperate in directing attention in space not only during the execution of goal-directed movements, but also in the absence of overt motor responses(46). In a Simon et al., 2002 (47) study participants performed various tasks, including attention, pointing, grabbing, saccades and calculating. Both precuneus and SPL were activated by saccadic and pointing task while theprecuneus showed more activation only for attention. Le et al., 1998 (48) reported that the shift of attention to visual stimuli, when compared to sustained attention, produced bilateral activation of precuneus and SPL.

Although, acurate recognition of complex emotions in facial expressions is usually more evident in the right hemisphere (42), in our expriments, sources were located in both or left hemispheres.

Both experiments were carried out with the purpose of contributing to the clarification of the open questions whether relatively early ERPs components can be modulate by facial expressions. We can conclude that N170 is modulated by expressions and it is possible to differentiate between expressions and neutral faces, but not between positive and negative expressions (happy and sad), which also holds true for the P250. Regarding the P100 we can conclude that it is not modulated by emotional expressions, in line with the notion that it is a quite early ERP type involved in low level processing. Our paradigm does therefore shed converging light on the neural correlates of the perception of emotional faces(4).

